# Behavioral and neural indices of affective coloring for neutral social stimuli

**DOI:** 10.1101/178384

**Authors:** Daniel W. Grupe, Stacey M. Schaefer, Regina C. Lapate, Andrew J. Schoen, Lauren K. Gresham, Jeanette A. Mumford, Richard J. Davidson

## Abstract

Emotional processing often continues beyond the presentation of emotionally evocative stimuli, which can result in affective biasing or coloring of subsequently encountered events. Here, we describe neural correlates of affective coloring and examine how individual differences in affective style impact the magnitude of affective coloring. We conducted functional magnetic resonance imaging in 117 adults who passively viewed negative, neutral, and positive pictures presented 2s prior to neutral faces. Brain responses to neutral faces were modulated by the valence of preceding pictures, with greater activation for faces following negative (vs. positive) pictures in the amygdala, dorsomedial and lateral prefrontal cortex, ventral visual cortices, posterior superior temporal sulcus, and angular gyrus. Three days after the MRI scan, participants rated their memory and liking of previously encountered neutral faces. Individuals higher in trait positive affect and emotional reappraisal rated faces as more likable when preceded by emotionally arousing (negative or positive) pictures. Additionally, greater amygdala responses to neutral faces preceded by positively valenced pictures were associated with greater memory for these faces three days later. Collectively, these results reveal individual differences in how emotions spill over onto the processing of unrelated social stimuli, resulting in persistent and affectively biased evaluations of such stimuli.

## Introduction

Emotional experiences are inherently “sticky”. Responses to affectively significant events (whether positive or negative) are rarely temporally constrained to the time of exposure, but can persist long after these events have passed (Frijda et al., 1991; Verduyn et al., 2015). Completely unrelated events, encountered in the wake of these evocative emotional experiences, may be processed through a particular affective filter, consequently “coloring” the perception or evaluation of unrelated events (Anderson, Siegel, White, & Barrett, 2012; Murphy & Zajonc, 1993). For example, harsh feedback from a graduate student’s supervisor may result in an intractable state of negative affect, which is subsequently reflected in overly critical evaluations of the statistics exams she grades that evening. On the other hand, after having a manuscript accepted in a prestigious journal, the same student may experience sustained positive affect that positively colors her evaluations of the next stack of exams. Notably, under certain circumstances such affective coloring has the potential to result in biased long-term evaluations of, or memory for, subsequent unrelated events or interactions (Lapate et al., 2017; Tambini et al., 2016). While recent studies have begun to illuminate the neural correlates of affective coloring (Lapate et al., 2017, 2016; Tambini et al., 2016), it is unknown whether and how inter-individual variability in neural mechanisms or trait-like indices of affective style bear on affective coloring outcomes.

Individual differences in the temporal dynamics of emotional responding, or “affective chronometry” (Davidson, 1998), may be critically important for the extent of such affective coloring. It is now well established that individual differences in affective chronometry are critical for both positive well-being (Davidson 2004; Heller et al. 2013; Schaefer et al. 2013) and risk of psychopathology (Blackford et al. 2009; Burke et al. 2005; Dichter and Tomarken 2008; Lapate et al. 2014; Larson et al. 2006; Schuyler et al. 2014; Siegle et al. 2002) in addition to negative physical health outcomes (Brosschot et al., 2006). Investigating affective coloring may afford insight into one pathway through which the temporal dynamics of emotion can impact psychological health: Individuals who quickly recover to a baseline state may exhibit minimal behavioral or biological indicators of spillover of emotion from one context to the next, whereas those who are slower to recover from negative events (or who are more capable of “savoring” positive events) will be more susceptible to this spillover or coloring. For example, previous studies utilizing daily diary methods have shown that while individuals with depression do not necessarily show greater reactivity to stressful life events, the impact of interpersonal stressors “spills over” into greater experiences of negative affect on subsequent days (Gunthert et al., 2007), and that the extent of this spillover predicts poorer response to cognitive behavioral therapy (Cohen et al., 2008). The laboratory investigation of individual differences in affective coloring may thus provide novel mechanistic insight into affective pathology and its treatment in a more carefully controlled setting.

To that end, our lab has developed an fMRI paradigm designed to measure the impact of well-characterized affective stimuli on immediate (neural) and long-term (behavioral) indices of affective coloring (specifically, the coloring of neutral social stimuli; Figure 1). In this task, participants passively view negative, neutral, and positive pictures that are followed by a face displaying a neutral expression, and are instructed to indicate the gender of the face (a low-level categorization task). To investigate affective coloring of neural responses, we compare brain responses to neutral faces as a function of the preceding picture valence. To investigate enduring behavioral indices of affective coloring, we present the same faces to participants 3 days later and test the extent to which memory or liking of these faces is modulated by the type of picture that initially preceded these faces.

**Figure 1.**
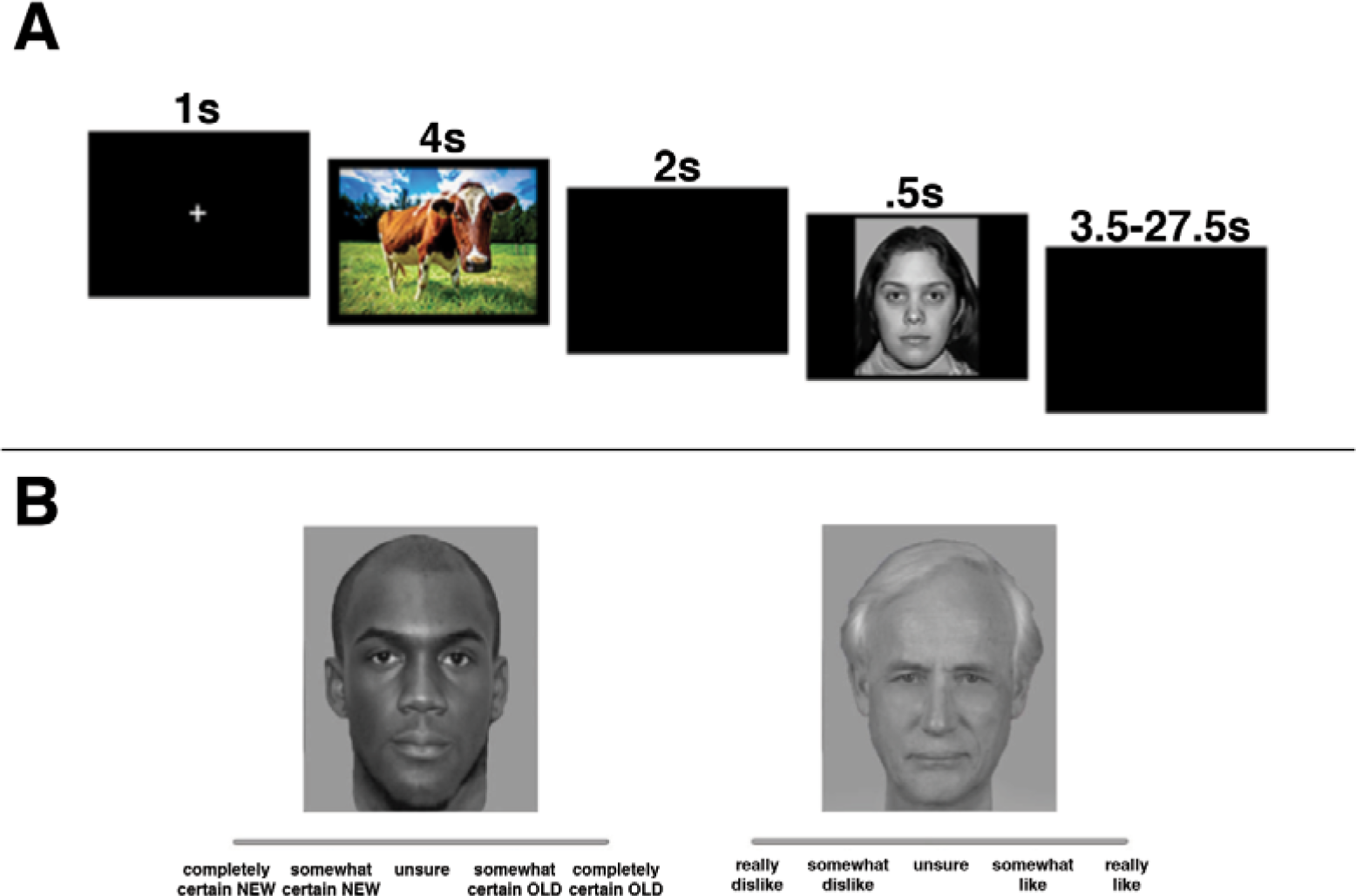
(A) Schematic of fMRI trial structure. On each trial, participants passively viewed a 4s picture from the International Affective Picture System (Lang et al., 2008). Following a 2s delay, a neutral face was presented and participants pressed a button to indicate the gender of the face. (B) Three days following the scan, participants used an online or paper rating scale to provide memory then likability ratings for 45 neutral faces viewed during the fMRI scan and 45 novel neutral faces.

## Method

### Participants

Participants for this study were drawn from Midlife in the United States (MIDUS), a national longitudinal study of health and well-being across the lifespan (http://www.midus.wisc.edu). For the current study, we included participants from the MIDUS “refresher” sample who were enrolled beginning in 2012 in an effort to expand and refresh the original MIDUS sample with younger age cohorts. The majority of the MIDUS refresher sample was recruited through random digit dialing of adults throughout the U.S. The refresher sample also includes an oversampling of African American participants from Milwaukee, WI who were recruited by door-to-door solicitation. Functional magnetic resonance imaging (fMRI) task data were collected on a subset of 122 MIDUS refresher participants living in the Midwest who were able to travel to our laboratory, including 35 participants from the Milwaukee subsample. All participants were right handed and were screened to ensure MRI compatibility, and all provided informed consent prior to all study procedures.

### MRI data collection

MRI data were collected on a 3T scanner (MR750 GE Healthcare, Waukesha, WI) using an 8-channel head coil. We collected 3 sets of echo planar images during the fMRI task (231 volumes, TR=2000, TE=20, flip angle=60°, field of view=220 mm, 96x64 matrix, 3-mm slice thickness with 1-mm gap, 40 interleaved sagittal slices, and ASSET parallel imaging with an acceleration factor of 2). A 3-dimensional magnetization-prepared rapid gradient-echo sequence (MPRAGE (Mugler and Brookeman, 1990)) was used to acquire a T1-weighted anatomical image for functional data registration (TR=8.2, TE=3.2, flip angle=12°, field of view=256 mm, 256x256 matrix, 160 axial slices, inversion time = 450 ms). The MRI session also included the acquisition of field map images, a resting-state scan, diffusion-weighted data, and perfusion data.

### Functional MRI task

During fMRI scanning, participants passively viewed pictures from the International Affective Picture System (IAPS; Lang, Bradley, and Cuthbert 2008), presented for 4s (Figure 1). Each of 3 runs contained 10 pictures each from negative, neutral, and positive valence categories, for a total of 90 picture presentations. Pictures were presented using a pseudorandom trial order with the requirement that no more than 2 pictures from the same valence category be presented in a row. Within each valence category, stimuli were randomly selected for each participant. Each picture was followed by a 2s inter-stimulus interval and a .5s neutral face presentation. Participants were instructed to press one of two buttons to indicate the gender of the face, a low-level categorization task chosen to confirm task engagement without interfering with natural processing of the faces or preceding IAPS pictures. Faces were followed by an inter-trial interval between 3.5-27.5s (mean duration=7.5s). A 1s crosshair appeared before the start of each trial to orient participants’ attention.

This trial timing was chosen based on simulations conducted using optseq2 (<u>https://surfer.nmr.mgh.harvard.edu/fswiki/optseq2</u>), with the constraint that faces be presented within 4s of picture presentation. This relatively brief ISI increased statistical power to observe a spillover effect by maximizing the number of trials within the scan session. Specifically, we compared model efficiency for trial timings that included 1) average ISIs of 2s and 3s, 2) jittered ISIs of 0s, 0.5s, and 1s, and 3) ITIs that varied (on average) between 4.5-8s. Based on these simulations, the fixed 2s interval provided the greatest power to detect effects of interest during the face period. Because this fixed ISI may raise concerns regarding collinearity of picture and face regressors, we calculated the variance inflation factor (VIF) for each regressor. A VIF cutoff of either 5 or 10 is typically used to indicate problematic levels of collinearity, and VIF values for the face regressors fell close to 5 (mean for picture regressors = 4.22-4.72; mean for face regressors = 4.82-5.45). Although the VIF values for face regressors in some cases exceeded this more conservative threshold, collinearity generally does not affect the Type I error rate, but rather can lead to more variable estimates and thus negatively impact statistical power (Mumford et al., 2015). In particular, more variable estimates at the level of individual subjects will average out across subjects, and should thus stabilize with a larger sample size. In contrast, collinearity in higher-level analyses (which was not the case in this study) may result in invalid inferences regarding the magnitude of effect sizes (for discussion, see Mumford et al., 2015).”

Average normative arousal ratings were equivalent for negative (5.46 ± 0.66) and positive (5.47 ± 0.53) IAPS pictures, which were both greater than those for neutral pictures (3.16 ± 0.42; *t*s > 16.1, *p*s < 0.001). Pictures within each valence category were matched on luminosity and visual complexity, and for the number of pictures rated to be social in content. Faces were drawn from the XM2VTSDB multi-modal face database (Messer et al., 1999), the NimStim database (Tottenham et al., 2009), and the Montreal Set of Facial Displays of Emotion (Beaupre et al., 2000). In light of our diverse subject population, these faces included equal proportions of male and females with a broad range of ages and ethnicities. Faces were cropped just above the hair and below the chin, converted to black and white, and edited (e.g., to remove distinctive facial hair and eyeglasses). A total of 45 neutral faces were presented, and each face was presented following 2 randomly selected pictures from the same valence category.

### FMRI data processing and analysis

FMRI data processing was carried out using FEAT (FMRI Expert Analysis Tool) Version 6.00, part of FSL (FMRIB’s Software Library, www.fmrib.ox.ac.uk/fsl). Preprocessing steps included removal of the first 4 volumes, motion correction using MCFLIRT, removal of non-brain regions using BET, spatial smoothing using a Gaussian kernel with 5mm full-width at half-maximum, grand-mean intensity normalization, and high-pass temporal filtering. We excluded 5 participants following first-level data processing (1 for excessive movement [>1mm mean framewise displacement across the entire run], 2 for failing to provide behavioral responses, and 2 for distorted functional data), resulting in a final sample size of 117 for fMRI analyses (64 female/53 male; Milwaukee subsample *N* = 35; mean age ± SD = 48.3 ± 11.8; range = 26-76). Additionally, due to excessive motion and/or failure to provide behavioral responses on at least 25% of trials, we excluded 1 run from 10 participants and 2 runs from 1 participant; for the other 106 participants, we included data from all 3 runs in group-level analyses.

The first-level general linear model included 6 regressors of interest, consisting of separate picture and face regressors for each picture valence (Negative/Neutral/Positive). Regressors of no interest included mean-centered response times to face presentations, 6 motion parameters with their first‐ and second-order derivatives, and a confound regressor for each volume with > .9 mm framewise displacement (Siegel et al., 2014). Additional processing steps included correction of time series autocorrelation, resampling of functional data to 2mm^3^ isotropic voxels, and registration to Montreal Neurological Institute template space.

As a complementary approach, we modeled amygdala time series data using a set of 7 finite impulse response (FIR) basis functions. This allowed for model-free estimation of the hemodynamic response from the time of picture onset through the first 14 seconds of each trial (corresponding to the minimum trial length). We then extracted empirically derived activation estimates for each trial type separately. For group-level analyses we averaged these estimates across individuals at each volume, calculated valence and arousal contrast estimates at each time point, and conducted paired-sample *t* tests for these contrasts.

### Group analysis of fMRI data

Primary group analyses for the fMRI data were conducted using two orthogonal contrasts for both the picture and face epochs: a “Valence” contrast (Negative vs. Positive) and an “Arousal” contrast (.5*(Negative+Positive)-Neutral). For significant effects involving the Arousal contrast, follow-up analyses examined whether effects were driven more by the Negative or Positive condition. For primary analyses focused on the amygdala, we extracted mean parameter estimates for each of these contrasts from voxels with ≥ 50% probability of being assigned to the bilateral amygdala from the ≥ Harvard-Oxford probabilistic atlas (volume=4040 mm^3^; Desikan et al., 2006; Figure 2a), which was chosen as an *a priori* ROI prior to data analysis. In addition to this *a priori* region of interest, we conducted whole-brain voxelwise analyses on the same 2 contrasts for the picture and face periods. Cluster threshold correction was applied using a Z threshold of 3.09 and a cluster-corrected significance value of *p* < 0.05.

**Figure 2.**
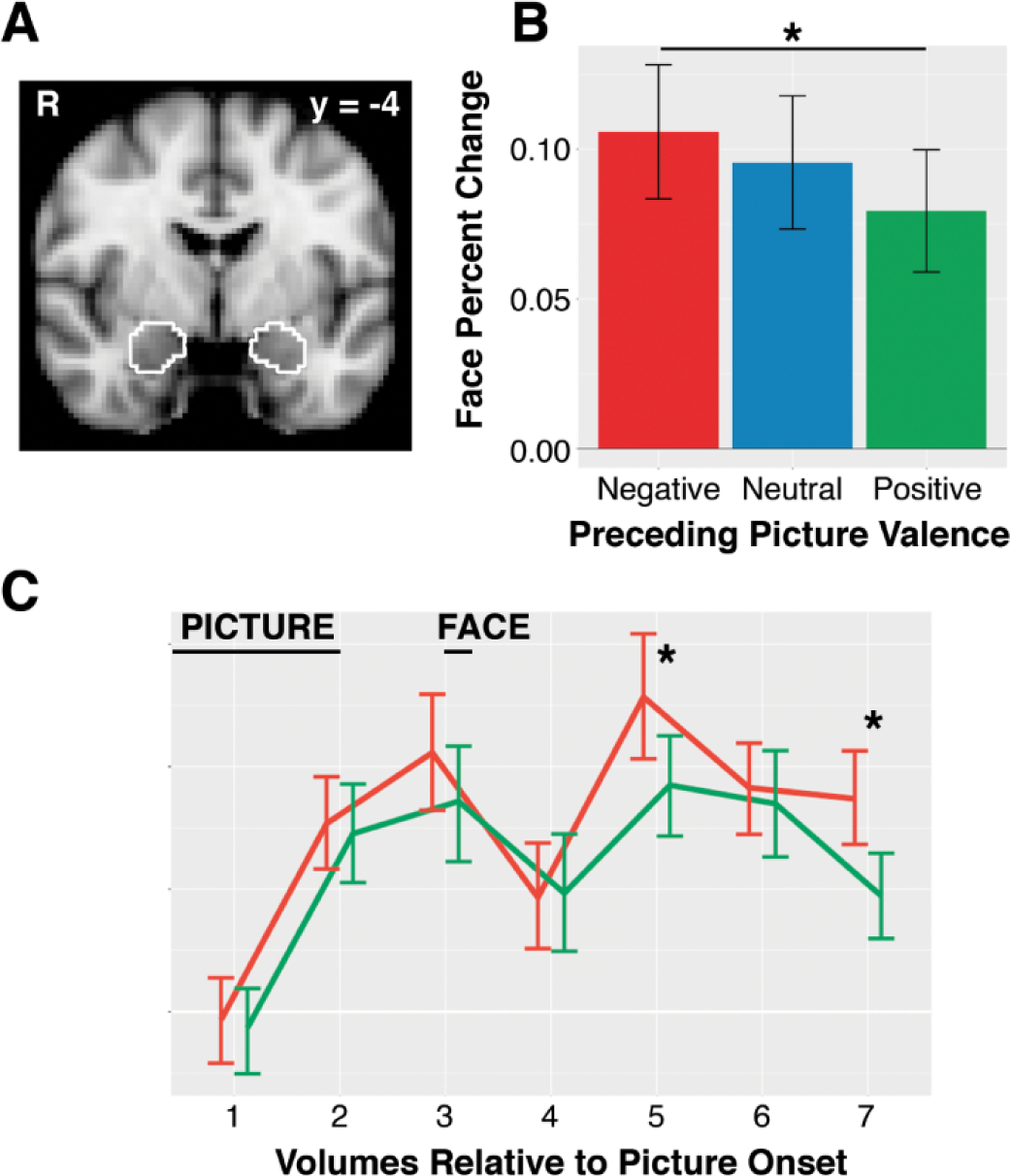
(A) Anatomically defined amygdala region of interest (ROI). (B) Mean percent signal change values across the amygdala ROI for neutral faces presented following negative, neutral, and positive pictures. (C) Estimated signal from the amygdala ROI using a set of finite impulse response basis functions revealed complementary evidence for greater amygdala activation to faces following negative vs. positive pictures. Notes: Error bars represent 95% confidence intervals. * *p* < 0.05.

### Behavioral data collection and analysis

Participants were asked to log onto the internet 3 days after the MRI session, and complete ratings of the 45 faces they viewed during the MRI task, which were interspersed with 45 novel faces. Each of these 90 faces was presented for an unlimited duration with one of two continuous rating scales that asked participants to rate their memory and liking of these faces. The memory scale had equidistant anchor points labeled “completely certain new”, “somewhat certain new”, “unsure”, “somewhat certain old”, and “completely certain old”; the liking scale had equidistant anchor points labeled “really dislike”, “somewhat dislike”, “unsure”, “somewhat like”, and “really like”. Memory ratings were always completed prior to liking ratings.

Memory and liking ratings were obtained from 105 and 104 participants, respectively. Of these participants, 16 were not comfortable using the online rating scale or did not have access to a computer; these individuals instead completed ratings using paper packets. Ratings collected electronically and on paper were both converted to values ranging between −1 and 1, with higher values indicating greater liking of, or greater confidence of having previously seen, each face. These ratings were completed exactly 3 days after the scan in 85 out of 89 subjects who completed ratings online (2 participants completed ratings 4 days after the scan, and 1 each completed ratings 7 and 9 days after the scan). All 16 participants completing paper ratings reported completing these exactly 3 days after the scan.

Analysis of these behavioral data tested the extent to which each participant’s long-term memory or liking of neutral faces was colored by the content of preceding IAPS pictures from the scan session. We conducted paired *t* tests on these behavioral data using the same contrasts as with the fMRI data in order to test whether the valence or arousal of pictures had a persistent impact on behavioral evaluations of subsequently presented faces. We predicted that 1) memory for neutral faces would be enhanced for faces following emotionally arousing pictures (Negative or Positive, relative to Neutral), and 2) liking of neutral faces would be enhanced for faces following Positive relative to Negative pictures.

### Individual differences analyses

We conducted analyses to explore the impact of individual differences in both affective style and amygdala activation on the affective coloring of behavior, as assayed by 3-day liking and memory ratings. We examined 4 self-report measures of affective style: state Positive and Negative Affect from the Positive and Negative Affect Schedule (Watson et al., 1988); the Reappraisal subscale of the Emotion Regulation Questionnaire (Gross and John, 2003); and total score on the Psychological Well-Being Scale (Ryff and Keyes, 1995). For each of these 4 indices we calculated Pearson correlation coefficients with subsequent ratings of face liking and memory, both as a function of preceding pictures’ Valence and Arousal. We corrected for multiple comparisons testing using a conservative Bonferroni-corrected *p* value of 0.003125 (4 measures * 2 behavioral ratings (liking/memory) * 2 contrasts (Valence/Arousal).

We also tested 2 specific hypotheses regarding relationships between individual differences in affectively-modulated amygdala responses to neutral faces and long-lasting behavioral evaluations of these faces. First, as an extension of findings that the amygdala mediates arousal-enhanced memory for emotional stimuli (Canli et al., 2000; Kensinger and Corkin, 2004), we hypothesized that amygdala responses to faces *following* arousing pictures would be associated with greater memory for those faces 3 days later. Second, we correlated amygdala responses to faces following Negative vs. Positive pictures (Valence contrast) with liking ratings, with the prediction that more negative affective coloring of faces (as manifest in stronger amygdala responses) would be associated with relatively lower levels of liking 3 days later. In addition to these *a priori* hypotheses, tested by calculating bivariate Pearson correlation coefficients, we also conducted exploratory whole-brain, voxelwise correlation analyses to identify other regions in which activation was associated with subsequent behavioral evaluations of neutral faces.

For each of these individual differences analyses, participants with behavioral data > 3 SD from the mean were excluded as univariate outliers. Additionally, for all effects identified as significant, the primary analysis (*t* test or correlation) was repeated as a linear regression model, controlling separately for effects of age, gender, or Sample (Milwaukee/main sample). Statistical maps for all whole-brain, voxelwise analyses are available at http://neurovault.org/collections/2803/.

## Results

### Amygdala responses to neutral faces are modulated by the valence of preceding pictures

During picture viewing, amygdala activation was modulated by emotionally arousing pictures ((.5*(Negative+Positive)>Neutral); *t*(116) = 5.94, *p* < 0.001, *d* = 0.55), and also by the valence of these pictures (Negative>Positive; *t*(116) = 3.33, *p* = 0.0012, *d* = 0.31). Whole-brain activation during picture viewing was generally consistent with previous studies investigating affective picture viewing (e.g., Lindquist et al., 2015). For example, the arousal contrast yielded activation in ventromedial and dorsomedial PFC, lateral orbitofrontal cortex and inferior frontal gyrus, amygdala, midbrain, precuneus, and occipital cortices; and deactivation in lateral frontal poles, precuneus, and lateral occipital cortex (<u>https://neurovault.org/images/52490/)</u>. Greater activation for negative vs. positive pictures was seen in lateral and inferior frontal gyri, midbrain, occipital cortices into the ventral visual stream, and cerebellum; greater activation for positive vs. negative pictures was observed in the middle frontal gyrus and midline regions, including medial PFC, posterior cingulate, precuneus, and cuneus (<u>https://neurovault.org/images/52489/)</u>.

Amygdala responses to neutral faces were not modulated by the arousal contrast for preceding pictures (*t*(116) = −0.33, *p* > 0.7). However, amygdala responses to neutral faces were modulated by the valence of preceding pictures, with greater activation when the preceding picture was negative vs. positive (*t*(116) = 2.39, *p* = 0.027, *d* = 0.22; Figure 2b). We also estimated amygdala responses directly from the data using a set of FIR basis functions (Figure 2c). This model-free approach revealed clearly distinct hemodynamic responses in the amygdala to the picture and face epochs, with peak responses 3 volumes (6s) after picture onset and 2-3 volumes (4-6s) after face onset. Consistent with our primary analysis, we observed greater amygdala activation for the negative vs. positive condition at volumes 5 (*t*(116) = 2.76, *p* = 0.0067, *d* = 0.27) and 7 (*t*(116) = 3.68, *p* < 0.001, *d* = 0.39), i.e., corresponding to 4 and 8 seconds, respectively, after face onset. Results of the whole-brain, voxelwise FIR analysis for all volumes are available at https://neurovault.org/collections/3154/.

Notably, when including age as a covariate, we no longer observed a significant group effect of Valence (*t*(112) = 1.24, *p* = 0.22). Indeed, age showed a marginally significant correlation with amygdala activation for the Valence contrast (*r*(115) = 0.17, *p* = 0.064), with greater (Negative>Positive) amygdala activation in older adults. To visualize this age effect, we conducted a *post hoc* median split of the sample into adults younger than 50 (*N* = 58) and older than 50 (*N* = 59). Whereas older adults had significant amygdala modulation by preceding picture valence (*t*(58) = 2.75, *p* = 0.008, *d* = 0.36), younger adults showed no statistical difference (*t*(57) = 0.79, *p* = 0.44, *d* = 0.10; Figure 3). Valence modulation of the amygdala was also reduced to non-significance when controlling for gender (*t*(115) = 1.62, *p* = 0.11), although there were no gender differences in amygdala signal (*t*(115) = −0.02, *p* = 0.98). The amygdala effect remained significant when controlling for Sample status (*t*(115) = 2.20, *p* = 0.03).

**Figure 3.**
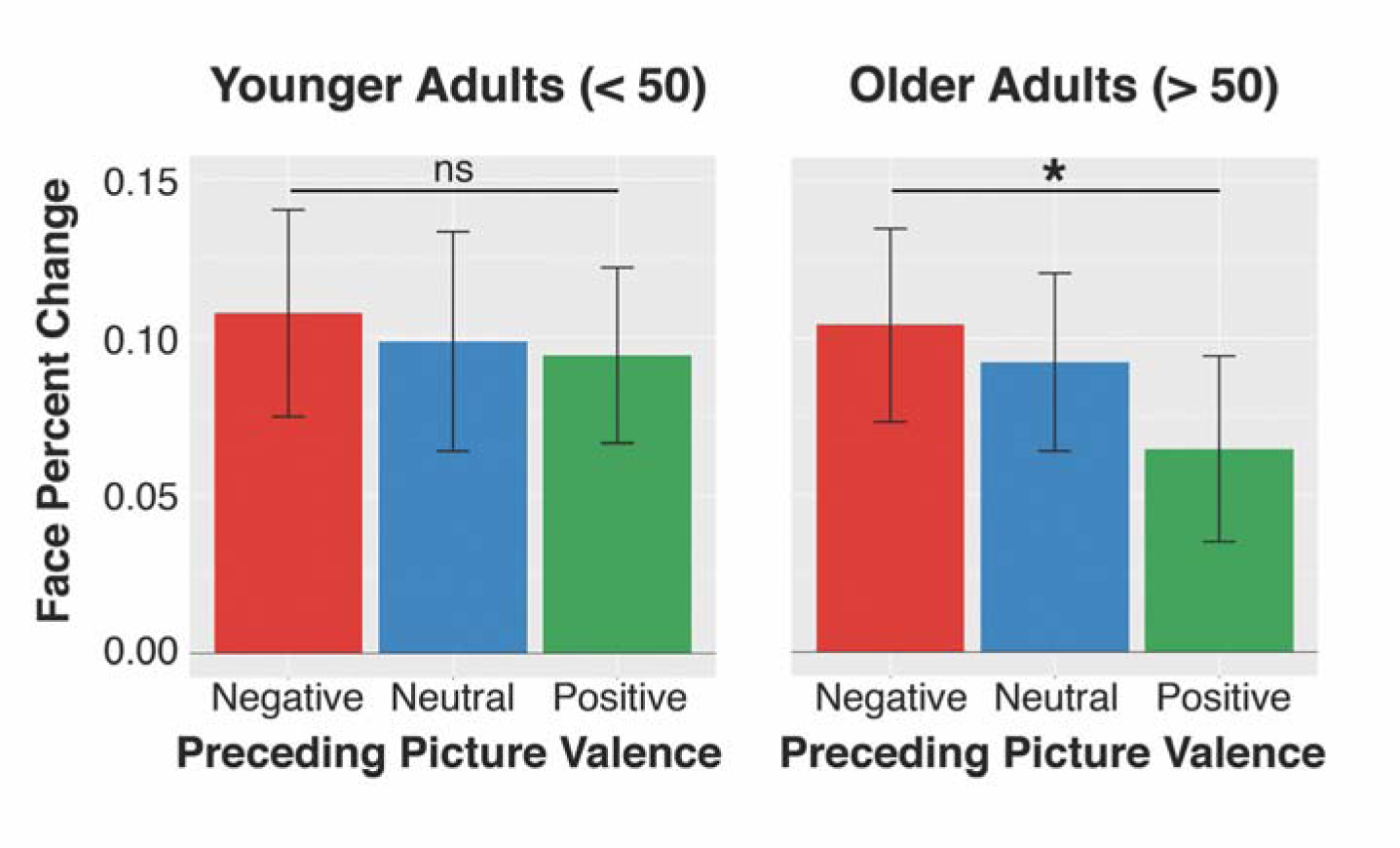
Mean percent signal change values across the amygdala ROI for neutral faces presented following negative, neutral, and positive pictures, for younger and older adults (based on a *post hoc* median split of the data). Older but not younger adults showed significant modulation of amygdala activation to faces based on the preceding valence of pictures, driven by a (non-significant) attenuation of amygdala activation to faces following positive pictures in older vs. younger adults (*t*(115) = 1.43, *p* = 0.16). Notes: Error bars represent 95% confidence intervals. * *p* < 0.01.

### Whole-brain responses to faces modulated by the valence of preceding pictures

We conducted whole-brain-corrected voxelwise searches to identify whether regions beyond the amygdala showed differential responses to neutral faces based on preceding pictures’ valence or arousal. For the valence contrast, we observed greater activation for neutral faces following negative vs. positive pictures in the dorsomedial prefrontal cortex (dmPFC), left inferior frontal gyrus, left cerebellum, and regions typically involved in face processing including lateral fusiform gyri bilaterally, right posterior superior temporal sulcus (STS), and bilateral angular gyri (Figure 4a and https://neurovault.org/images/52492/). The arousal contrast revealed enhanced activation in occipital cortex for faces following emotionally arousing (negative or positive vs. neutral) pictures (Figure 4b and https://neurovault.org/images/52494/). All results held when controlling for covariates of age, gender, or Sample, and no regions were identified that showed effects in the opposite directions.

**Figure 4.**
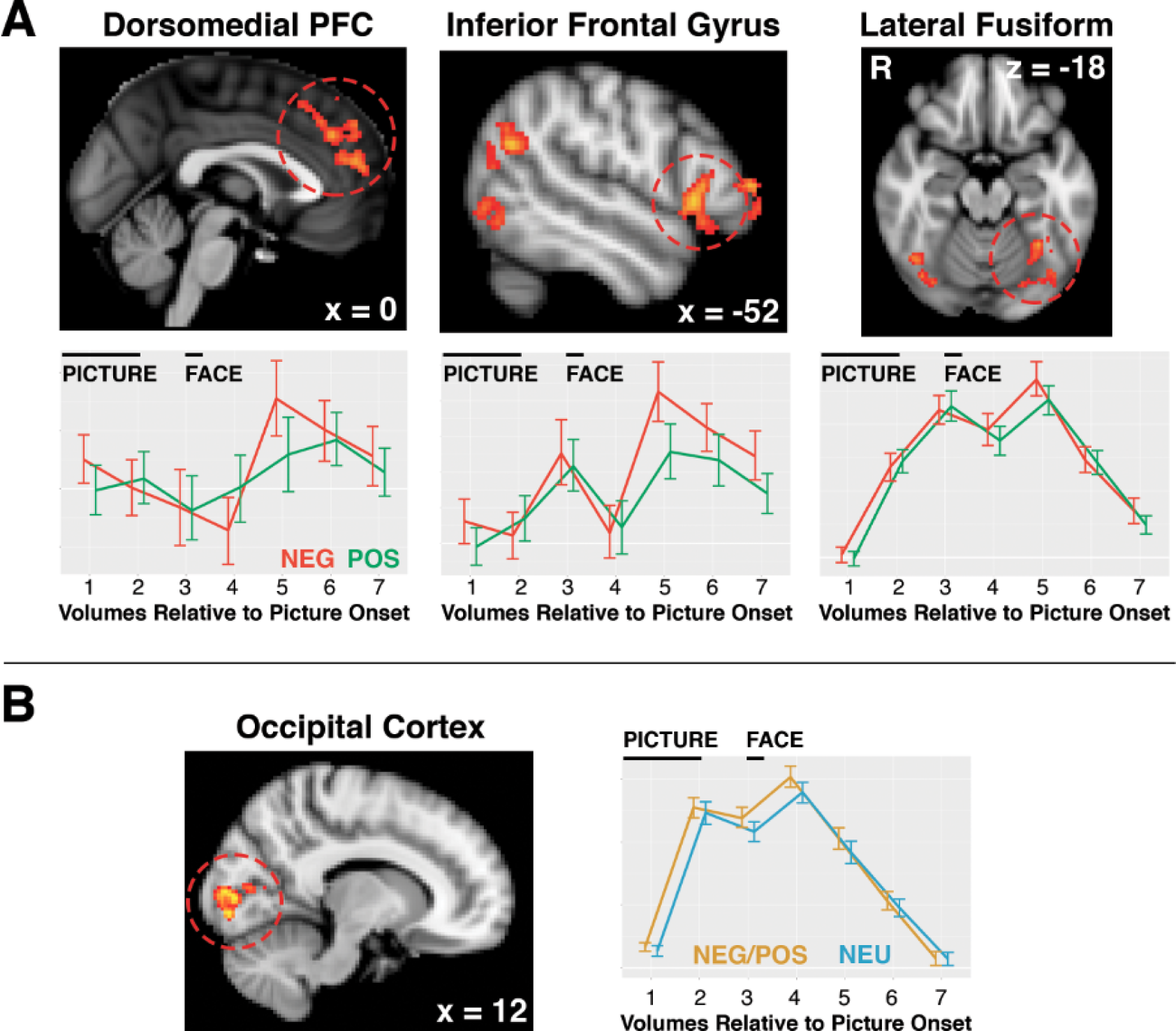
(A) Whole-brain voxelwise analysis for the Valence contrast revealed greater activation to neutral faces following negative vs. positive pictures in the dorsomedial prefrontal cortex, left inferior frontal gyrus, and lateral fusiform gyri bilaterally, in addition to the right posterior superior temporal sulcus, bilateral angular gyri, and left cerebellum. Activation estimates from a set of finite impulse response basis functions are presented for these functionally defined clusters for illustrative purposes only. (B) Whole-brain voxelwise analysis for the Arousal contrast revealed greater activation to neutral faces following negative *and* positive (relative to neutral) pictures only in the occipital cortex. Maps are presented at *p* < 0.05, using cluster threshold correction. Full statistical maps can be viewed at http://neurovault.org/collections/2803/.

### Individual differences in affective style and liking of neutral faces three days later

On an average of 3 days after the fMRI session, participants provided memory and then liking ratings of the 45 faces viewed during the fMRI session, intermixed with 45 face foils. Participants were able to successfully discriminate previously seen from novel foil faces (*t*(104) = 12.48, *p* < 0.001, *d* = 1.22), and they rated previously seen faces as more likable than novel faces (Zajonc, 1968; *t*(103) = 7.33, *p* < 0.001, *d* = 0.72). Contrary to hypotheses, there were no main effects of preceding pictures’ valence or arousal on long-term memory or liking ratings (|*t*s| < 0.9, *p*s > 0.4).

We next asked whether individual differences in self-report measures of affective style were associated with long-lasting affective coloring of neutral faces. Faces preceded by emotionally arousing (vs. neutral) pictures were rated 3 days later as more likeable by individuals higher in trait positive affect (*r*(100) = 0.25, *p* = 0.012, uncorrected; Figure 5a) and emotional reappraisal (*r*(99) = 0.34, *p =* < 0.001, Bonferroni-corrected α <0.05; Figure 5b). The relationship between reappraisal and liking still survived Bonferroni correction (α <0.05) when controlling for age, gender, or Sample. The relationship with positive affect remained significant (not correcting for multiple comparisons) when controlling for age, gender, or Sample (*t*s > 2.3, *p*s < 0.03). Positive affect and reappraisal were significantly correlated with one another (*r*(113) = 0.23, *p* = 0.012), and only reappraisal accounted for unique variance in face liking in a simultaneous regression model (reappraisal: *t*(98) = 3.01, *p* = 0.004; positive affect: *t*(99) = 1.73, *p* = 0.09).

**Figure 5.**
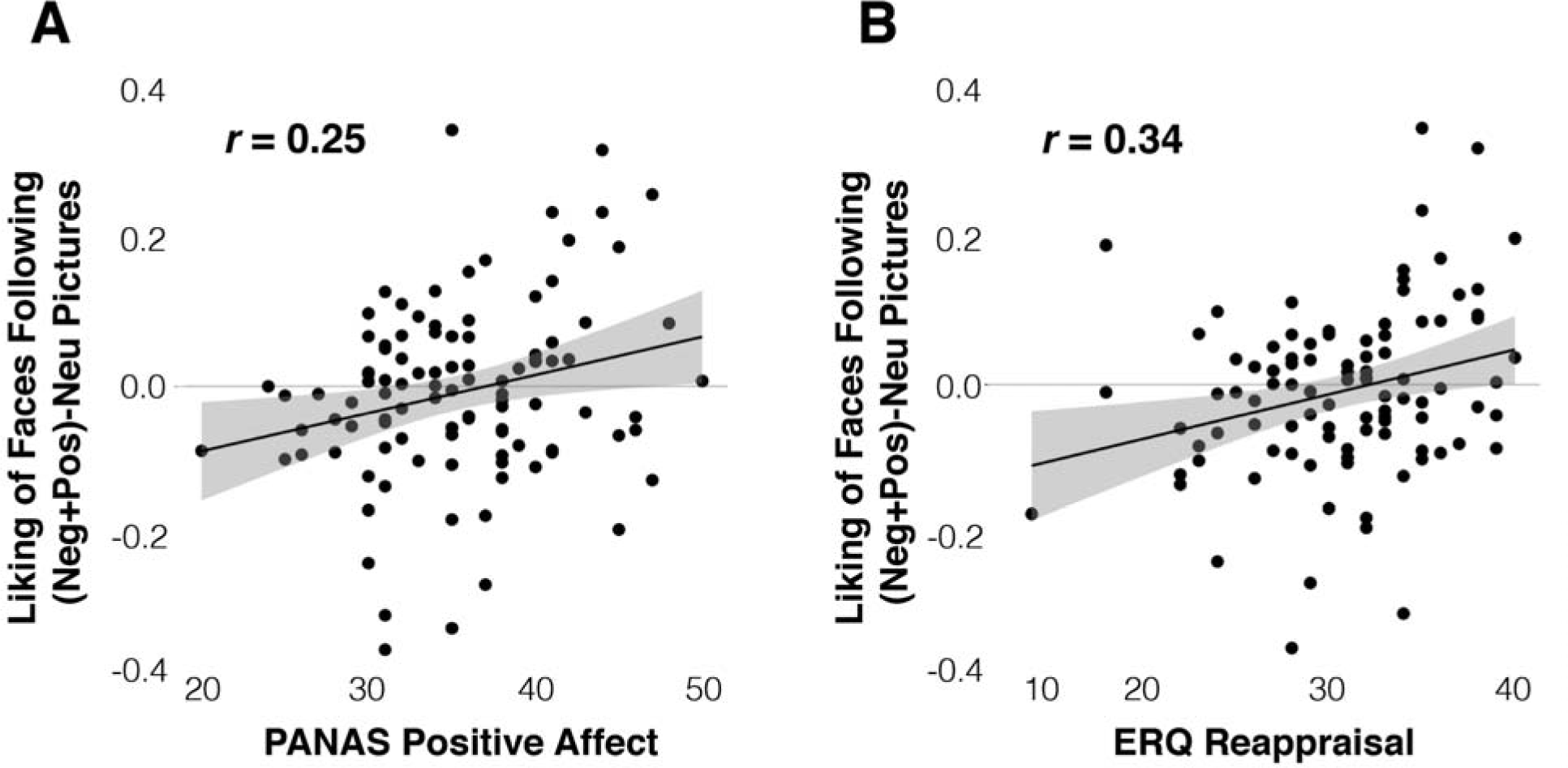
(A) Individuals higher in trait positive affect on the Positive and Negative Affect Schedule (PANAS) showed greater arousal-modulated liking of faces viewed during the MRI session, such that faces initially preceded by negative *or* positive (relative to neutral) faces were rated as more likable when viewed 3 days later. (B) A similar relationship was observed for the Reappraisal subscale of the Emotion Regulation Questionnaire (ERQ).

Follow-up analyses showed that these effects were present for faces following both positive and negative (relative to neutral) pictures (positive affect: *r*s > 0.20, *p*s < 0.05; reappraisal: *r*s > 0.25, *p*s < 0.02). No correlations were observed between liking ratings and trait negative affect or psychological well-being, or between memory ratings and any of these 4 self-report measures (|*r*s| < 0.14, *p*s > 0.1).

### Individual differences in amygdala activation and memory for neutral faces three days later

Consistent with predictions, individuals with increased amygdala activation to faces presented following emotionally arousing pictures had greater memory for those same faces when tested 3 days later (*r*(100) = 0.21, *p* = 0.039; Figure 6a). Planned follow-up analyses showed that this effect was driven by faces following positive vs. neutral pictures (*r*(99) = 0.21, *p* = 0.036; Figure 6b), with no relationship for faces following negative vs. neutral pictures (*r*(100) = −0.03, *p* = 0.78). The confidence interval for the difference between these correlations included zero (−0.01, 0.48), indicating that the relationships between positive and negative conditions were not significantly different (Zou, 2007). Amygdala responses to faces following positive pictures were associated with enhanced memory for those faces when controlling for age, gender, or Sample (*t*s > 2.1, *p*s < 0.04). In contrast, there was no relationship between differential amygdala responses to faces following negative vs. positive pictures and subsequent behavioral evaluations (liking or memory) of those faces 3 days later (*r*s| < 0.11, *p*s > 0.3). Exploratory analyses did not reveal any other brain regions in which affectively colored activation to neutral faces were associated with face ratings 3 days later (<u>https://neurovault.org/images/52495/</u> and https://neurovault.org/images/52496/).

**Figure 6.**
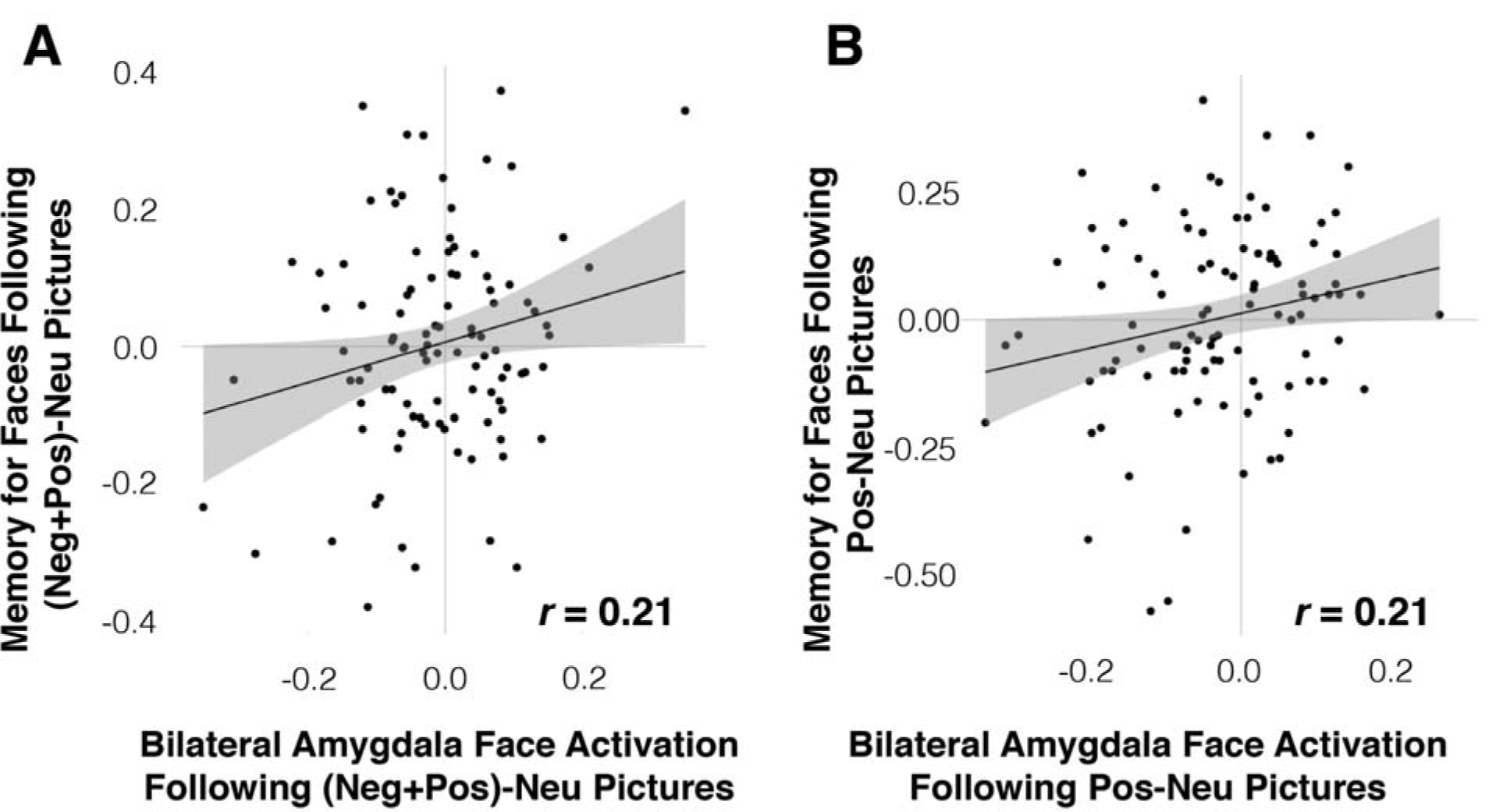
(A) Individuals with stronger amygdala responses to neutral faces following negative or positive (relative to neutral) pictures showed arousal-enhanced memory for neutral faces 3 days after the MRI session. (B) This effect was specific to the positive valence condition, as amygdala responses for faces following positive (vs. neutral) but not negative pictures predicted greater memory for these same faces 3 days later.

Collectively, these results show that short-term (neural) response to neutral faces are affectively colored by preceding emotional pictures, and that long-term (behavioral) indices of affective coloring vary as a function of individual differences in trait affective style as well as emotionally modulated amygdala activation.

## Discussion

In a large and diverse sample of healthy adults, we have demonstrated an important role for individual differences in emotional responding on determining enduring behavioral effects of affective coloring. We found that self-report measures of positive affective style were associated with more positive appraisals of neutral faces that 3 days earlier had been presented in the wake of either positive or negative (relative to neutral) pictures. We also found that greater amygdala responsivity for neutral faces appearing after emotional pictures was associated with increased memory for these faces 3 days later, an effect that somewhat unexpectedly was present for positive but not negative affective stimuli. Collectively, our results underscore the critical role of individual differences in determining how emotions spill over onto unrelated social stimuli, thus coloring our subsequent preferences and memory for these stimuli.

Contrary to predictions, we observed no main effects of emotional pictures’ valence or arousal on behavioral evaluations of the following neutral faces, assayed 3 days after the fMRI scan, across the sample. Instead, our results indicate that certain individuals are more prone than others to the influence of emotion stimuli on subsequent coloring. We found that individual differences in questionnaire measures of positive affective style (emotional reappraisal and trait positive affect) were robustly related to arousal-modulated liking of neutral faces viewed 3 days after the fMRI scan. For individuals high in these dimensions, the arousal elicited by pictures may be appraised in a more positive manner due to differences in baseline emotional traits (Schachter and Singer, 1962), and may thus have differential consequences for how other stimuli in the environment are processed – in this case, leading individuals to form persistent positive appraisals of neutral faces encountered in the wake of emotionally arousing pictures.

Neurally, greater amygdala responses to neutral faces following more arousing pictures (in particular, positive pictures) were associated with increased memory for those faces when viewed three days later. This result provides a novel contribution to studies on the role of the amygdala in emotional memory, which have previously shown that amygdala activation at encoding mediates the relationship between emotionally arousing stimuli and increased memory encoding or retrieval for those stimuli (Cahill et al., 1996; Canli et al., 2000; Kensinger and Corkin, 2004; LaBar and Phelps, 1998). Here, we demonstrate that memory for *neutral* faces, presented shortly after positive affective stimuli, is enhanced specifically in individuals who show elevated amygdala responses to those neutral faces (see also Tambini, Rimmele, Phelps, & Davachi, 2016). It is notable that emotionally modulated amygdala activation was linked to greater memory for positive and not for negative stimuli. Similarly, an earlier report found memory enhancement for neutral stimuli presented *prior to* positive stimuli, as opposed to a retrograde amnesic effect for neutral stimuli presented before negative stimuli (Hurlemann et al., 2005). Indeed, emotionally enhanced memory of neutral stimuli is more frequently observed when these neutral stimuli precede rather than follow emotional stimuli (Knight and Mather, 2009).

Notably, we did not identify a main effect of memory enhancement or impairment for emotional relative to neutral stimuli; again, as with findings related to affective style, this memory effect emerged purely as a function of our individual differences analytical approach. If replicated and extended in other samples, this finding may have important real-world consequences: those individuals who show prolonged amygdala activation in the immediate aftermath of emotionally arousing events may be more apt to process unrelated (but temporally proximal) neutral stimuli in the environment in a way that enhances encoding or later retrieval. Research into the temporal dynamics of emotional responses may thus bear on our understanding of emotional memory and its disruption in conditions such as posttraumatic stress disorder (Hayes et al., 2011) or depression (Burt, Zembar, & Niederehe, 1995; Mather et al., 2006). Depression in particular is increasingly linked to deficiencies in the experience of, or ability to sustain, positive affect (Heller et al., 2013, 2009; Kovacs et al., 2016). Based on our results, it is important to consider the possibility that altered affective coloring in psychopathology may not reflect excessive coloring by negative events, but rather deficient coloring by positive events, or the absence of what speculatively may be a protective “positivity bias” (cf. Alloy and Abramson, 1979).

In addition to the long-term behavioral consequences of affective coloring, our experimental paradigm allows for the examination of short-term neural correlates of affective coloring. We provide a novel theoretical contribution by demonstrating that amygdala responses to neutral social stimuli were differentially modulated by the valence of preceding emotional pictures, with greater responses following negative vs. positive pictures. Results from our model-free approach corroborated those of primary analyses, and provided visual evidence that this effect resulted from differential increases in amygdala activation following neutral face onset (as opposed to a sustained response to emotional pictures themselves). This coloring of amygdala responses was observed over a very short timescale, with neutral faces appearing only 2s after picture offset. It would be informative to probe the system at later time points to track the persistence of this effect, and to vary the picture-face ISI to probe individual differences in this persistence.

This amygdala effect was no longer significant when controlling for age, due to a (non-significant) increase in amygdala activation to faces following negative-positive pictures with increasing age. These findings may seem inconsistent with the literature on the positivity effect, which refers to biased information processing in older adults for positive vs. negative information. The positivity effect is consistently reflected in increased attention or memory for positive vs. negative information (Reed et al., 2014); importantly, the relationship we observed between amygdala responses to faces following positive pictures and subsequent memory for these faces remained significant while controlling for age. In addition to biased information processing, aging is also associated with relatively greater amygdala activation for positive vs. negative stimuli (Mather, 2016). This literature has focused primarily on amygdala activation *during* the processing of affective stimuli, and does not speak to post-stimulus activation or the temporal dynamics of amygdala responding. Indeed, in a study using corrugator EMG to assess affective chronometry, we previously identified slower recovery from negative IAPS stimuli in older women relative to younger women and men (van Reekum et al., 2011). This underscores the possibility that the relationship between aging and differential responses to positive vs. negative information may be qualitatively different *during* the processing of emotional stimuli vs. *after* stimulus processing (and for stimuli that appear during recovery from emotional stimuli), a possibility that should be more fully explored in future work.

Left lateral PFC and dorsomedial PFC regions also showed enhanced activation to neutral faces following negative vs. positive pictures. These same regions are consistently engaged during the voluntary regulation of emotional responses (Buhle et al., 2013, Figure 1), although in the current study participants were not given explicit emotion regulatory instructions. Notably, the magnitude of these regions’ inverse coupling with the amygdala was recently linked to short-term reductions in affective coloring behavior in the aftermath of fearful face viewing (Lapate et al., 2016). Moreover, these regions show significant age-related declines in cortical thickness (Fjell and Walhovd, 2010) and associated cognitive function (MacPherson et al., 2002). Further testing is warranted to explore whether age-associated changes in the structure and functionality of these prefrontal regions may underlie the pronounced affective coloring effect observed in the amygdala of older participants.

Similar valence-modulated activation of neutral faces was observed in the lateral fusiform gyrus and posterior STS, which compose a “core system” for face processing (Haxby et al., 2000). Previous studies have demonstrated that these regions are involved in representing distinct facial expressions of emotion (Fox et al., 2009; Harry et al., 2013; Vuilleumier and Pourtois, 2007), thought to reflect the downstream influence of emotional processing conducted in the amygdala (Haxby and Gobbini, 2011; Vuilleumier et al., 2004). The current findings support and extend these previous studies and theoretical model, suggesting that affective information may spill over from preceding and unrelated emotional events in a way that can color the perception or evaluation of unrelated social stimuli. Future work should explore the real-word impact of differential engagement of this social-affective neural circuitry with regard to biased social interactions or other behavioral outcomes.

In summary, we have presented a novel fMRI paradigm that reveals the impact of affective pictures on subsequent neural responses to neutral social stimuli, and uncovers individual differences in affective style and amygdala activation that are related to enduring behavioral indices of affective coloring. In light of relationships between trait-like differences in affective style and risk for developing psychopathology (Fredrickson et al., 2003; Shackman et al., 2016; Troy et al., 2010), and alterations in emotional memory and amygdala activation in conditions such as depression and posttraumatic stress disorder (Hayes et al., 2011; Mather et al., 2006), these results suggest that an affective coloring framework may provide novel insight into neural and behavioral mechanisms of affective psychopathology. This work paves the way for subsequent investigations in affective disorders marked by alterations in affective chronometry, such as truncated experiences of positive affect and slower recovery from negative challenges, which will allow us to relate these temporal dynamic differences to behavioral indices of emotional coloring and different symptom measures.

## Author contributions

S.M. Schaefer, R.C. Lapate, and R.J. Davidson developed the study concept and contributed to study design. Testing and data collection was performed by A.J. Schoen and L.K. Gresham. D.W. Grupe, S.M. Schaefer, R.C. Lapate, and J.A. Mumford performed the data analysis and interpretation. D.W. Grupe drafted the manuscript, and all authors provided critical revisions and approved the final version of the manuscript for submission.

## Acknowledgements

The authors thank Luke Hinsenkamp, Ben Hushek, Ayla Kruis, Phoebe Marquardt, Karthik Aroor, Nathan Vack, Michael Anderle, Ron Fisher, and Scott Mikkelson for assistance with data collection, processing, and analysis.

